# Structural determinants of REMORIN nanodomain formation in anionic membranes

**DOI:** 10.1101/2022.08.16.501454

**Authors:** Anthony Legrand, Daniel G. Cava, Marie-Dominique Jolivet, Marion Decossas, Olivier Lambert, Vincent Bayle, Yvon Jaillais, Antoine Loquet, Véronique Germain, Marie Boudsocq, Birgit Habenstein, Marisela Vélez, Sébastien Mongrand

**Affiliations:** Laboratoire de Biogenèse Membranaire, CNRS, Univ. Bordeaux, (UMR 5200), F-33140 Villenave d’Ornon, France; Laboratorio de Biofuncionalización de Superficies (Biofunctional Surfaces) Instituto de Catálisis y Petroleoquímica CSIC c/Marie Curie 2. Cantoblanco 28049, Madrid, Spain; Univ. Bordeaux, CNRS, Bordeaux INP, CBMN, UMR 5248, IECB, Pessac, France; Laboratoire Reproduction et Développement des Plantes, Université de Lyon, ENS de Lyon, UCB Lyon 1, CNRS, INRA, F-69342 Lyon, France; Institute of Plant Sciences Paris-Saclay (IPS2), Université Paris-Saclay, CNRS, INRAE, Université Evry, Université Paris-Cité, 91190 Gif-sur-Yvette, France

## Abstract

Remorins are a family of multigenic phosphoproteins of the plasma membrane, involved in biotic and abiotic plant interaction mechanisms, partnering in molecular signaling cascades. Signaling activity of remorins depends on their phosphorylation states and subsequent clustering into nano-sized membrane domains. The presence of a coiled-coil domain and a C-terminal domain is crucial to anchor remorins to negatively charged membrane domains, however the exact role of the N-terminal intrinsically disordered domain (IDD) on protein clustering and lipid interactions is largely unknown. Here we combine chemical biology and imaging approaches to study the partitioning of group 1 remorin into anionic model membranes mimicking the inner leaflet of the plant plasma membrane. Using reconstituted membranes containing a mix of saturated and unsaturated PhosphatidylCholine (PC), PhosphatidylInositol Phosphates (PIPs), and sterol, we investigate the clustering of remorins to the membrane and monitor the formation of nano-sized membrane domains. REM1.3 promoted membrane nanodomain organization on the exposed external leaflet of both spherical lipid vesicles and flat supported lipid bilayers. Our results reveal that REM1.3 drives a mechanism allowing lipid reorganization, leading to the formation of remorin-enriched nanodomains. Phosphorylation of the N-terminal IDD by the calcium protein kinase CPK3 influences this clustering and can lead to the formation of smaller and more disperse domains. Our work reveals the phosphate-dependent involvement of the N-terminal IDD in the remorin-membrane interaction process by driving structural rearrangements at lipid-water interfaces.

**Summary heading:** Using reconstituted membranes, we demonstrated the clustering of the plant protein remorins StREM1.3 to the lipid bilayer external leaflet and monitor the formation of nanodomains of the protein.

## Introduction

Membrane nanodomain compartmentation is a common mean to organize and regulate activities at the plasma membrane (PM) (Jaillais and Ott, 2020). Failure to do so may result in critical dysfunction, as exemplified for small GTPases, such as the proto-oncogene Ras (Prior et al., 2012; Zhou et al., 2018) or the yeast Rho GTPases Cdc42 (Sartorel et al., 2018). Studying the mechanisms underlying such an organization appears crucial in order to understand how membrane compartmentation impacts biological function. In this regard, a plethora of studies were performed on various PM nanodomain systems including both lipids and proteins. For example, in animals, Ras nanodomains (Weise et al., 2011; Zhou et al., 2017), ganglioside GM1 nanodomains (Ewers et al., 2010; Yuan and Johnston, 2001) and caveolae (Parton and Simons, 2007) are prominent targets of intense biochemical and biophysical studies. Unicellular organisms such as yeasts or bacteria also organize their PMs into nanodomains (Bramkamp and Lopez, 2015; Spira et al., 2012). Reconstitution of membrane nanodomain systems furthers our understanding of their underlying molecular mechanism and provides a formidable platform for *in vitro* biophysical studies (Cebecauer et al., 2018; Mouritsen and Bagatolli, 2015; Simons and Vaz, 2004). These studies highlight the importance of lipid species, particularly cholesterol and anionic phospholipids, involved in direct protein-lipid interactions. Rho-related GTPase of Plants (ROP) are Rho GTPases, which are capable of forming nanoclusters in the inner leaflet of the plant PM. Single-molecule super-resolution microscopy revealed that ROP6 is stabilized by PhosphatidylSerine (PS) into PM nanodomains, which are required for auxin signaling (Platre et al., 2019; Smokvarska et al., 2020).

Another well-known protein that clusters in nanodomains of the inner leaflet of the plant PM belongs to the multigenic family of proteins called remorins (REMs) (Raffaele et al., 2007). REMs encompass a broad range of functions from protection against abiotic stress and host-pathogen biotic interaction, to inception of symbiosis and hormone signaling response (Gouguet et al., 2020). REMs belong to a family of 6 phylogenic groups with distinct N-terminal regions (Raffaele et al., 2007), which can label a plethora of nanodomains of different size, shape and localization (Jarsch et al., 2014). REMs are bound to the PM’s inner leaflet through an unconventional mode of targeting through the C-terminal peptide called REM-CA (REM-C-terminal Anchor, Figure 1A). The inner leaflet of the plant PM, which is facing the cytosol, is highly negatively charged due to the presence of acidic phospholipids, such as PS, as well as PhosphatidylInositol Phosphates (PIPs), notably PhosphatidylInositol 4-Phosphate (PI4P) that accumulate in the PM (Barbosa et al., 2016; Platre et al., 2018, 2019; Simon et al., 2016; Synek et al., 2021). It is worth noting that PIPs can form nanodomains on their own (Bilkova et al., 2017; Furt et al., 2010; Ji et al., 2015; Motegi et al., 2021), and polycationic peptides induces the clustering of PIPs (van den Bogaart et al., 2011).

**Figure 1.**
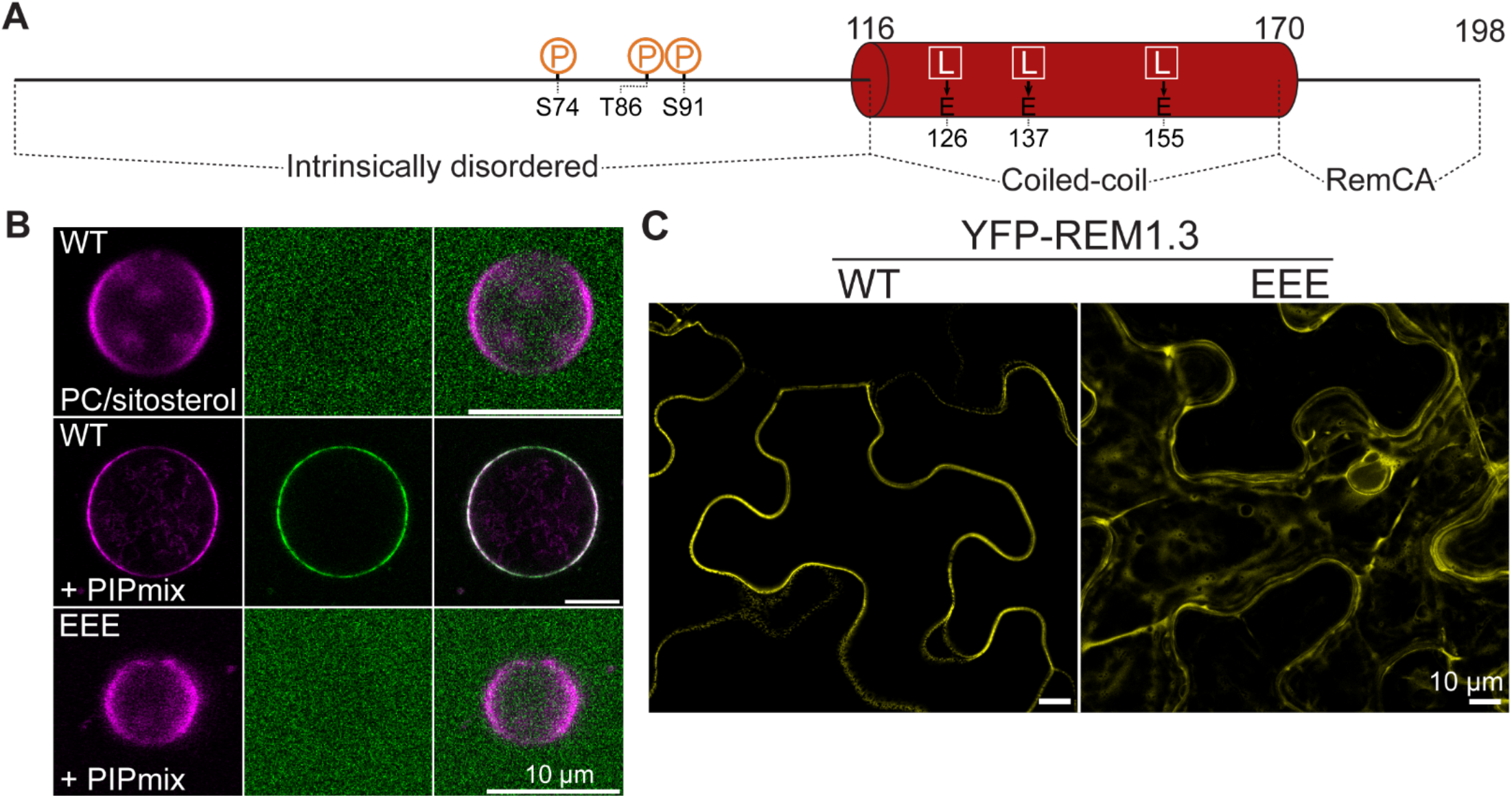
Minimal setting to address REM1.3’s organization in artificial membranes. Disrupting of REM1.3’s coiled-coil domain hampers membrane targeting. (A) Stick figure representing the different domains of StREM1.3. Red cylinder: coiled-coil domain with critical hydrophobic residues (white) and point-mutations (black) used in this study. P represents phosphorylation sites. REM-CA: Remorin C-terminal anchor. (B) Membrane targeting of GFP-REM1.3_86-198_ in GUVs composed of DPPC/DLPC/sitosterol, 8/50/42/mol% (top) or against DPPC/DLPC/sitosterol/PIPmix 50/26/8/16 mol% (middle) or GFP-REM1.3_86-198_^EEE^ against GUVs of DPPC/DLPC/sitosterol/ PIPmix 50/26/8/16 mol% (bottom) (molar percentages). *Left*: RhodPE channel (magenta). *Middle*: GFP-REM1.3_86-198_ channel (green). *Right*: composite images from both channels. Observation statistics are given in Table S1. (C) Membrane targeting of GFP-REM1.3 WT or EEE transiently expressed in *N. benthamiana* leaves. Scale bars: 10 μm.

The potato (*Solanum tuberosum*) group 1 isoform 3 REM, further called REM1.3, was particularly studied because its REM-CA is not palmitoylated (no cysteine residues is present). Therefore, the anchoring is entirely due to protein/lipid and protein/protein interaction, without the need for any post-translational modification of a lipid moiety to the REM-CA. Biophysical and mutagenesis studies of this REM-CA domain displayed a complicated interaction mechanism with the PM, based on a balance between: (1) electrostatic interactions through Lysines and Arginines and PI4P; (2) hydrophobic effects between REM-CA inside the lipid bilayer; (3) anchor stabilization by phytosterols (Gronnier et al., 2017; Konrad et al., 2014; Legrand et al., 2019; Perraki et al., 2012; Raffaele et al., 2009). Full length REM1.3 was shown to segregate into liquid-ordered (lo) lipid domains both *in vitro* by solid state NMR (Legrand et al., 2019), and *in vivo* using an environmental sensitive fluorescent probe (Gronnier et al., 2017). Our groups further evidenced the relevant role of REM1.3’s coiled-coil domain, stabilized by three hydrophobic contacts led by leucine residues leading to trimerization (Figure 1A). Indeed, coiled-coil-disruptive single mutations from leucines to prolines resulted in a strong defect in PM targeting (Martinez et al., 2018). Finally, REMs also contain an N-terminal intrinsically disordered domain (IDD, Figure 1A) that can be phosphorylated (Marin et al., 2012; Marín and Ott, 2012; Perraki et al., 2018). Importantly, this IDD is not involved in membrane anchoring *per se* (Legrand et al., 2019), but is likely important in protein-protein interactions as described for many IDDs (Bah et al., 2015; Uversky, 2013). For example, we showed that the IDD of group 1 REMs can be phosphorylated by a Calcium Protein Kinase isoform 3 (CPK3), and those phosphorylations induced nanodomain rearrangement with a significant increase of the mean square displacement of EOS-REM1.3 *in vivo* and a partial loss of nanodomain stability (Perraki et al., 2018).

In this article, we present an *in vitro* reconstitution of a plant REM nanodomain, in a minimal synthetic membrane mimicking the PM’s inner leaflet, in order to understand the mutual role of lipids, coiled-coil and phosphorylation status of the IDD. We yield a pictorial view of the localization process of REM1.3 in membranes up to a single molecule level by high-resolution imaging using confocal laser scanning, Cryo-Electron Microscopy (CryoEM) and tapping-mode Atomic force Microscopy (AFM). Partitioning of REM1.3 into this newly designed anionic model membrane of a nanodomain system addressed three biologically relevant questions: (1) what is the minimal set of partners required to make REM1.3 nanodomains; (2) what is the contribution of lipids and the REM1.3 protein in nanodomain formation; (3) how phosphorylation of REM1.3 modifies nanodomain organization? We postulate a molecular mechanism where the nanodomain formation would be REM1.3-driven and PIP- and sitosterol-dependent. In this view, phosphorylation of REM1.3 by CPK3 would modify protein-protein interactions, disrupting nanodomain organization into smaller and more disperse domains, thus phenocopying our *in vivo* observations by Single-Particle Tracking-Photoactivated Localization Microscopy (Gronnier et al., 2017).

## Material and methods

### Material and plant growth conditions

All lipids come from Avanti with the exception of phosphoinositides mix (PIPmix) from bovine brain (P6023, Sigma-Aldrich). GIPCs were purified from cauliflower, as described in (Mamode Cassim et al., 2021). For transient expression, *Nicotiana benthamiana* plants were cultivated in controlled conditions (16 hours photoperiod, 25°C). The GFP-StREM1.3 *Arabidopsis thaliana* expressing line was crossed with the *pss1-3+/-*mutant (Platre et al., 2018). The resulting F1 progeny was grown on sulfadiazine to select *pss1-3+/-*seedling and these plants were then self-fertilized. In F2 generation, *pss1-3-/-*seedlings were selected on the basis of their agravitropic phenotype (Platre et al., 2019) and analyzed by confocal microscopy. GFP-positive plants were selected and the localization of GFP-StREM1.3 marker was analyzed and quantified. F3 seeds are kept heterozygous for the *pss1-3* mutation.

### Protein production and purification

REM1.3_86-198_ and REM1.3 were produced and purified as described in (Legrand et al., 2019). GFP-REM1.3_86-198 WT_ and GFP-REM1.3_86-198,EEE_ were constructed as follows, from N-to C-terminal: GFP (Zacharias, 2002) – linker (LESTSPWKKAGS) – REM1.3_86-198_ (WT or L125E/L137E/L155E (EEE)). The corresponding DNA sequences were ordered from Eurofins Genomics and cloned in pET-24a between NdeI and XhoI. They were produced in BL21-DE3-pLys cells in LB medium with 30 μg/mL kanamycin. At OD_600_ = 0.6, 1 mM IPTG was used to induce protein expression at 18°C overnight. Cells were pelleted at 6000 g for 20 min at 4°C, resuspended in lysis buffer (20 mM HEPES 150 mM NaCl 20 mM imidazole 1 mM PMSF 0.02% NaN_3_ pH = 7.4 with Complete protease inhibitor cocktail, Roche) and sonicated on ice. The lysate was centrifuged at 15557 g for 30 min at 4°C and the supernatant was loaded on a HisTrap column (GE Healthcare) equilibrated in 20 mM HEPES 150 mM NaCl 20 mM imidazole 0.02% NaN_3_ pH = 7.4). GFP-REM1.3_86-198_ was eluted with elution buffer (20 mM HEPES 150 mM NaCl 400 mM imidazole 0.02% NaN_3_ pH = 7.4), dialyzed against 10 mM HEPES 150 mM NaCl 0.02% NaN_3_ pH 7.4) at 4°C overnight, which triggered protein aggregation (Martinez et al., 2018). The turbid protein sample was centrifuged at 100000 g for 2 h at 4°C, and the pellet was discarded. After two more rounds of centrifugation, the last supernatant was kept and contained non-aggregated pure GFP-REM1.3_86-198_. Protein concentration was assessed by absorbance at 280 nm (ε_280_ = 41400 M^-1^.cm^-1^ according to Expasy ProtParam). All proteins were stored at 4°C.

Recombinant CPK3-GST was produced in BL21-DE3-pLys and purified as previously reported (Boudsocq et al., 2012).

### Giant unilamellar vesicles (GUVs) formation

Lipids at 10 g/L were mixed in organic solvent. About 20 μL were spread on Teflon disks, which were individually stored in small beakers. They were dried for at least 1h under vacuum with desiccator. Using a bubbler, lipids were pre-hydrated under a stream of N_2_-saturated H_2_O for 20 min. About 5 mL of 300 mM sucrose was gently layered on top of the disk (enough volume to cover it fully). From this point, care was taken not to shake the beaker to avoid breaking nascent GUVs. After a 34°C of overnight incubation, GUVs were collected using a severed pipette tip (to avoid shearing) and stored at 4°C until further use. GUVs were stable for 1 week.

### Fluorescence microscopy on Giant Unilamelar Vesicles

Teflon-coated 50 μL observation chambers were coated with 5% BSA for 20 min at room temperature then washed three times with 10 mM 150 mM NaCl pH = 7.4. Using a severed P200 pipette tip, a drop of the GUV suspension was deposited, followed by about ≈ 1.8.10^−12^ moles of GFP-tagged protein if necessary. A slightly elevated cover slide was installed using double-tape face. Observations were carried out through optical oil, on a Zeiss LSM 880 confocal laser scanning microscopy system (Leica, Wetzlar, Germany) equipped with Argon, DPSS and He-Ne lasers and a hybrid detector. GFP was excited at 488 nm and dioleoyl-Rhodamine-PhosphoatidylEthanolamine (RhodPE) was excited at 565 nm.

### Plant cells live imaging

Live imaging was performed using a Zeiss LSM 880 confocal laser scanning microscopy system. YFP and GFP fluorescence were observed with excitation wavelengths of 488 nm and emission wavelengths of 490 to 550 nm. Subcellular localization of YFP-REM1.3_WT_ and YFP-REM1.3_EEE_ was observed 48 hours after infiltration of 3-week-old *N. benthamiana* leaves with *Agrobacterium tumefasciens* GV3101 carrying the appropriate constructs (OD_600nm_ = 0.2). For imaging of *Arabidopsis thaliana* root seedlings, plants were grown vertically for 6 days on a half strength MS plate in controlled conditions (16 hours photoperiod, 22°C). The PM Spatial Clustering Index (SCI) was calculated based on images taken from epidermal cells of the elongation zone of 6-day-old root seedlings. In order to quantify, experiments were performed using identical confocal acquisition parameters. The SCI was calculated as previously described in (Gronnier et al., 2017). Briefly, 10 μm lines were plotted across the samples and the SCI was calculated by dividing the mean of the 5% highest values by the mean of 5% lowest values. Three lines were randomly plotted per cell.

### CryoEM

Lipids were hydrated to 10 g/L with 10 mM HEPES 150 mM NaCl 0.02% NaN_3_ pH = 7.4, submitted to five freeze-thaw-vortex cycles (liquid N_2_, 42°C water bath), mixed with GFP-REM1.3_86-198_ to final concentrations of 0.5 g/L lipids and 0.9 μM GFP-REM1.3_86-198_, then incubated at room temperature for 1h. Samples were loaded on a glow-discharged holey carbon-coated copper 300 mesh grids, blotted with a filter paper and frozen in liquid ethane using an EM-GP plunge freezer (Leica). Images were acquired on a Tecnai F20 electron microscope (ThermoFisher Scientific) operated at 200 kV using an Eagle 4k_4k camera (ThermoFisher Scientific).

### Atomic Force Microscopy (AFM)

Lipids were mixed in CHCl_3_ (or CHCl_3_/MeOH (2/1 vol/vol) for PIPmix), dried under a gentle flow of N_2_, resuspended in 10 mM HEPES, 10 mM NaCl pH = 7.4 buffer to 1-2 mg/mL at 40°C, and extruded 25 times at a temperature of 40°C by passing the solution through a polycarbonate membrane of 0.05 μm pore size (Avanti® Mini-Extruder). A liposome solution was incubated on a mica surface at 30°C overnight in a sealed Petri dish at high humidity, where liposomes fused together to form a homogeneous supported lipid bilayer. Such bilayers remain homogeneous for 1-2h at room temperature (20°C). For heterogeneous supported lipid bilayers, incubation was performed at room temperature.

The mica surface was thoroughly rinsed with water to remove excess liposomes. First, pure lipid bilayers were imaged. Then, REM1.3 or REM1.3_86-198_ was added in a 250/1 lipid/protein molar ratio and allowed to incubate for 30 min. The surface was then rinsed with buffer to remove unbound protein before imaging. Images were taken with an AFM from Agilent, model 5500, in tapping mode. The AFM tips used were specific for soft tapping, gold coated with a spring constant of 0.28 N/m.

### *In vitro* REM phosphorylation assay

The *in vitro* kinase assay was performed as previously described (Perraki et al., 2018). Briefly, CPK3-GST (WT or D202) was incubated with 1-2 μg of 6His-REM1.3 in the kinase reaction buffer (20 mM Tris HCl pH 7.5, 10 mM MgCl_2_, 1 mM DTT, 50 μM cold ATP, ATP [γ-33P] 2 μCi per reaction, 1 mM CaCl_2_) for 30 min at RT. The reaction was stopped with Laemmli buffer. Protein samples were heated at 95 °C for 3 min and separated by SDS-PAGE on 12% acrylamide gel. After migration, the gel was dried before exposing against a phosphorScreen to reveal radioactivity on a Typhoon Imaging system (GE Heathcare). The gel was then rehydrated for Coomassie staining.

## Results

### Anchoring of REM1.3 to lipid vesicles mimicking the inner leaflet of a plant plasma membrane

A first step towards reconstituting nanodomains of REM1.3 *in vitro* was to produce Giant Unilamellar Vesicles (GUVs) containing both saturated and unsaturated Phosphatidylcholine (PC) (namely DiPalmitoyl-PC (DPPC) and DiLinoleoyl-PC (DLPC)) and 8 mol% β-sitosterol, a well-established lipid mixture to promote lipid phase separation into liquid-ordered (lo) and liquid-disordered (ld) domains at room temperature (Cacas et al., 2016; Furt et al., 2010). In addition, GUVs were also supplemented with 16 mol% of a mix of PhosphoInositides (PIPs) and PhosphatidylSerine (PS) (called PIPmix, containing 50/20/15/15 mol% of PS/PI/PIP/PIP_2_ (Gronnier et al., 2017)). These lipids are the major anionic lipids constituting the PM’s inner leaflet to which REM1.3 is expected to bind (Gronnier et al., 2017; Legrand et al., 2019). To assess protein binding, we used a truncated GFP-tagged REM1.3, GFP-REM1.3_86-198_ (Figure S1), lacking its N-terminal intrinsically disordered domain (IDD) so that only the minimal anchoring machinery remains (Figure 2A). Images were taken by confocal microscopy after addition of GFP-REM1.3_86-198_ to the GUVs. As expected, GFP-REM1.3_86-198_ decorated 77% of the GUVs containing the PIP/PS lipid mixture (Table S1), but not those containing only PC and sitosterol, with which the GFP fluorescence signal remained in the buffer (Figure 1B). RhodPE labelling was non-uniform, indicating a partial phase separation (Figure 1B), even in absence of protein. This is expected for a complex mixture of high- and low-phase transition temperature PCs, i.e. DPPC and DLPC, sitosterol and anionic lipids, i.e. POPS and PIPs (Feigenson and Buboltz, 2001; Ingólfsson et al., 2014; Salvemini et al., 2014). Likewise, GFP-REM1.3_86-198_ labelling of liposomes was heterogeneous, partially mimicking, although with a much broader size, what can be observed in vivo for REM1.3 at the PM (Figure 3A,D).

**Figure 2.**
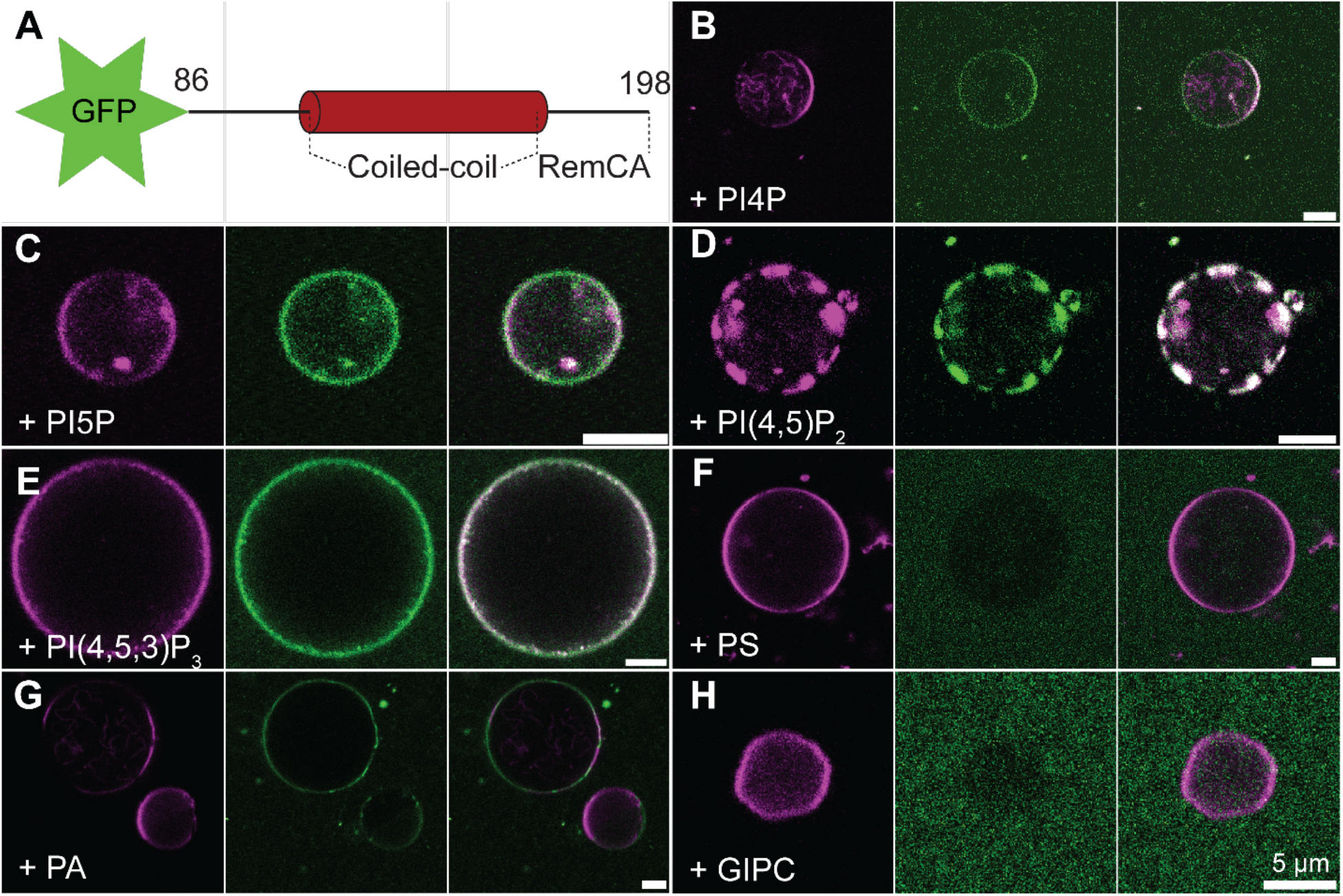
REM1.3 binds to GUVs containing PIPs or PA, but neither PS, nor GIPCs. (A) Stick figure of the purified protein GFP-REM1.3_86-198_ used. GUVs containing DPPC/DLPC/sitosterol 50/26/8 mol% supplemented with 16 mol% POPI4P (B), POPI5P (C), DOPI(4,5)P_2_ (D), PAPI(3,4,5)P3 (E), POPS (F), POPA (G), or GIPC (H), were mixed with ≈ 1.8.10^−12^ moles of GFP-REM1.3_86-198_. *Left*: RhodPE channel (magenta). *Middle*: GFP-REM1.3_86-198_ channel (green). *Right*: composite images from both channels. Scale bars: 5 μm. Observation statistics are given in Table S1.

**Figure 3.**
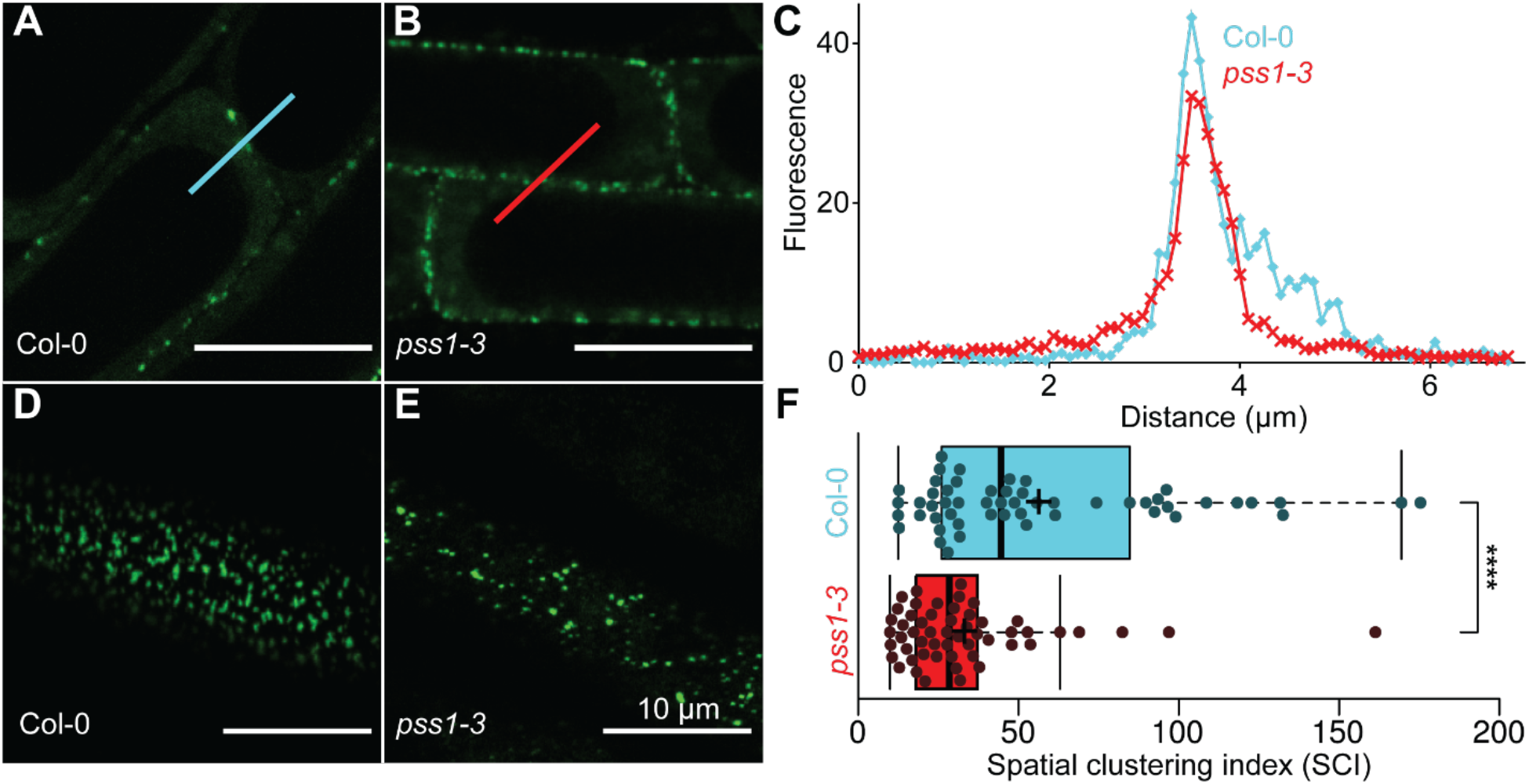
PS defines nanoclustering pattern, but not membrane targeting of REM1.3 *in vivo*. (A,B) Cross section confocal images of epidermal cells of Col-0 (A) or *pss1-3* KO (B) *Arabidopsis thaliana* 6-day-old root seedlings. (C) Fluorescence profiles for traces in (A, cyan) and (B, red) are drawn. (D,E) Surface view confocal images of epidermal cells of Col-0 (D) or *pss1-3* KO (E) *A. thaliana* 6-day-old root seedlings. (F) Spatial clustering index (SCI) was calculated using surface view confocal images (D,E). Tukey boxplots show the mean fluorescence intensity and the SCI of YFP-REM1.3 of at least 18 cells from 3 independent experiments. Significance was determined using a Student’s t-test (****: p<0.001). Scale bars: 10 μm.

### Coiled-coil homo-oligomerization of REM1.3 is essential for PM anchoring

We previously showed that REM1.3’s coiled-coil domain was crucial for membrane targeting *in vivo* (Martinez et al., 2018). Targeting was drastically hampered by mutation of its three conserved hydrophobic core residues L126, L137 and L155 to prolines. However, prolines are also likely to disrupt helices. We wished to inquire whether the disruption of the helical fold, or the lack of oligomerization, was responsible for this behavior. In order to maintain helical integrity, we mutated L126, L137 and L155 to negatively charged glutamates (REM1.3_EEE_) (Figure 1A). Confocal microscopy showed that GFP-REM1.3_86-198,EEE_ could only bind to 6% of all observed GUVs *in vitro* and most of the GFP fluorescent signal stayed in the buffer (Figure 1B). We further confirmed *in vivo* that full length YFP-REM1.3_EEE_ was unable to attach to the PM and stayed in the cytosol, when transiently expressed in *Nicotiana benthamiana* epidermal leaves (Figure 1C).

Thus, both the REM-CA and the coiled-coil domains are necessary and sufficient to target the protein to the PIPmix-containing lipid bilayer. This experimental setup allowed us to further characterize the lipid binding specificity of REM1.3.

### REM1.3 mainly targets anionic lipids with exposed phosphate groups

As a first step to address the role of lipids in the binding of REM1.3, we assess which lipids are important in this process. According to the literature (Gronnier et al., 2017; Legrand et al., 2019; Perraki et al., 2012), we formulated the hypothesis that GFP-REM1.3_86-198_ (Figure 2A) preferentially binds to any lipid with a surface-exposed phosphate. To test this hypothesis, we generated liposomes with the same PC/sterol mixture supplemented with different individual PIP species with one to three free phosphate groups at 16 mol% i.e. Palmitoyl-Oleoyl-PI4P (PO-PI4P), Palmitoyl-Oleoyl-PI5P (PO-PI5P), Di-Oleoyl-PI(4,5)P_2_ (DO-PI(4,5)P_2_) and Palmitoyl-Arachidonic-PI(3,4,5)P_3_ (PA-PI(3,4,5)P_3_). Figure 2B-E and Table S1 showed that GFP-REM1.3_86-198_ could also bind to all of these anionic phospholipids.

We assessed whether PS, another abundant anionic lipid of the inner leaflet of plant PM, allowed the binding of GFP-REM1.3_86-198_ to liposomes. We generated liposomes with 16 mol% of Palmitoyl-Oleyl-PS (POPS), but GFP-REM1.3_86-198_ was not at all able to bind to those GUVs (Figure 2F), unless a much higher percentage of POPS (32 mol%) was used (Figure S2, Table S1). However, it was able to bind to GUVs containing 16 mol% of Palmitoyl-Oleoyl-Phosphatidic Acid (POPA), albeit with a low binding frequency of 14% (Figure 2G, Table S1). Thus, the affinity of GFP-REM1.3_86-198_ for PS seems much lower compared to its affinity for phospholipids with exposed phosphate groups like PIPs and PA.

As a final control, we tested glycosyl phosphatidylinositol ceramide (GIPC), an abundant anionic sphingolipid of the plant PM’s present in the outer leaflet of the plant PM (Cacas et al., 2016; Cassim et al., 2020). Yet, GFP-REM1.3_86-198_ did not bind to DLPC/sitosterol/GIPC 1/1/1/ GUVs (Figure 2H, Table S1).

### PS is not necessary for REM1.3 nanodomain formation *in vivo*

We further tested the apparent superfluity of PS for REM1.3’s membrane targeting *in vivo* by taking advantage of the *Arabidopsis thaliana* mutant lines of the PS synthase (*pss1-3*) completely lacking PS (Platre et al., 2018). GFP-REM1.3 was expressed in wild-type (WT) and *pss1* mutant background. Confocal fluorescence microscopy on *Arabidopsis thaliana* root seedlings showed GFP-REM1.3 was still targeted to the PM in the absence of PS. Furthermore, GFP-REM1.3 was partially organized in nanodomains at the PM of the *pss1-3* mutant. Thus, PS is dispensable for the addressing of REM1.3 to PM nanodomains (Figure 3A-E). However, the nanodomains appeared larger and more diffuse in *pss1-3* than in the WT, as measured by the Spatial Clustering Index (SCI) (Figure 3F), suggesting that PS may indirectly participate in the spatial organization of REM1.3 *in vivo*.

### Cryo-EM observations showed that REM1.3 is organized periodically into nanodomains on lipid vesicles

To observe individual oligomers of GFP-REM1.3_86-198_ at a nanometric scale, we needed a much higher resolution than what conventional confocal microscopy can provide. We thus adapted our previously described sample preparation protocol for CryoEM, replacing GUVs with large unilamellar vesicles (LUVs) and increasing the PIPmix content from 16 to 24 mol%, to increase the probability to spot protein binding events. When necessary, 0.9 μM of GFP-REM1.3_86-198_ were added and the mixture was incubated at room temperature for 1h before freezing.

Our protocol allowed us to obtain smooth, protein-free lipid vesicles (Figure 4A). In the presence of GFP-REM1.3_86-198,_ we observed patterned dots decorating the vesicle’s outer leaflets likely corresponding to a patch of proteins, while their inner leaflets remained smooth (Figure 4B). These protein patches shared many expected nanodomain features: (1) a finite domain size, here about 100 nanometers along the equatorial plane of the liposome; (2) a local enrichment in GFP-REM1.3_86-198_ organized along the membrane with center-to-center distance between dots of about 6.6 ± 1.1 nm; (3) a local thickening of the outer leaflet of membrane by ≈ 5 nm, due to the presence of GFP-REM1.3_86-198_ (Figure 4C).

**Figure 4.**
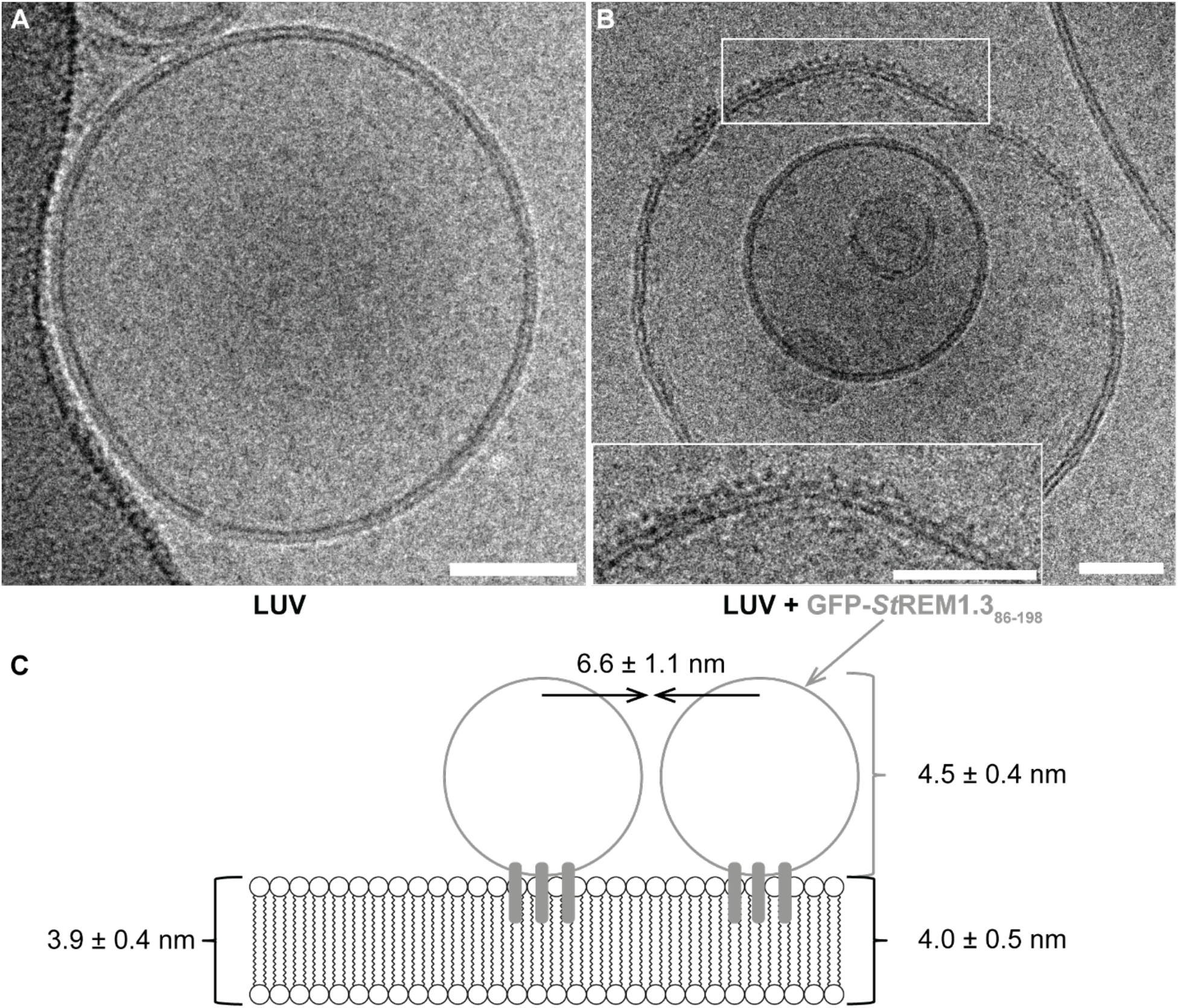
Cryo-EM reveals nanodomain segregation of GFP-REM1.3_86-198_ in LUVs containing PIPmix. (A) A protein-free Large Unilamellar Vesicles (LUV) composed of DPPC/DLPC/sitosterol/PIPmix (50/18/8/24 mol%). (B) GFP-REM1.3_86-198_ bound to a LUV. Notice the uneven distribution of protein and total thickness along its equatorial plane. Insert: zoom onto a GFP-REM1.3_86-198_-rich membrane domain. (C) Schematic representation of three GFP-REM1.3_86-198_ bound to a membrane: the 3 grey lines represent REM-CA, the balls represent the 3 REM-IDD-coiled-coil fused to GFP. Measured thicknesses for membrane, membrane under REM1.3 and REM1.3-REM1.3 are provided (n=20). Scale bars: 50 nm.

This data demonstrates that we are indeed able to reconstitute *in vitro* membrane nanodomains of GFP-REM1.3_86-198_ in lipid vesicles.

### The formation of REM1.3 membrane nanodomains is protein-driven and the shape and size of domains are phosphorylation-dependent

To gather more detailed insights into the REM1.3’s nanoclustering mechanism, we used Atomic Force Microscopy (AFM) on supported lipid bilayers to monitor the behavior of REM1.3 when attached to the membrane. We first started from a “smooth”, homogeneous lipid bilayer composed of DPPC/DLPC/sitosterol/PIPmix generated by heating the lipid mixture to remove pre-existing lipid phase separation. In absence of protein, no differences in height were observed on the surface of the supported lipid bilayer (Figure 5A left). The addition of the truncated REM1.3_86-198_ (purified without the GFP tag, see Figure S1) triggered the formation of thread-like domains about 8-10 nm thicker than the bulk membrane (Figure 5A, middle). To question the role of β-sitosterol, we performed AFM experiments in absence of β-sitosterol in the saturated-unsaturated PC mixture. Figure S3 clearly showed that the absence of β-sitosterol prevented REM1.3_86-198_ to increase the thickness of the membrane. Therefore, the presence of anionic lipids (Figure 1) and phytosterol (Figure S3) is not only required for the REM1.3 anchoring, but also for domain formation *in vitro*.

**Figure 5.**
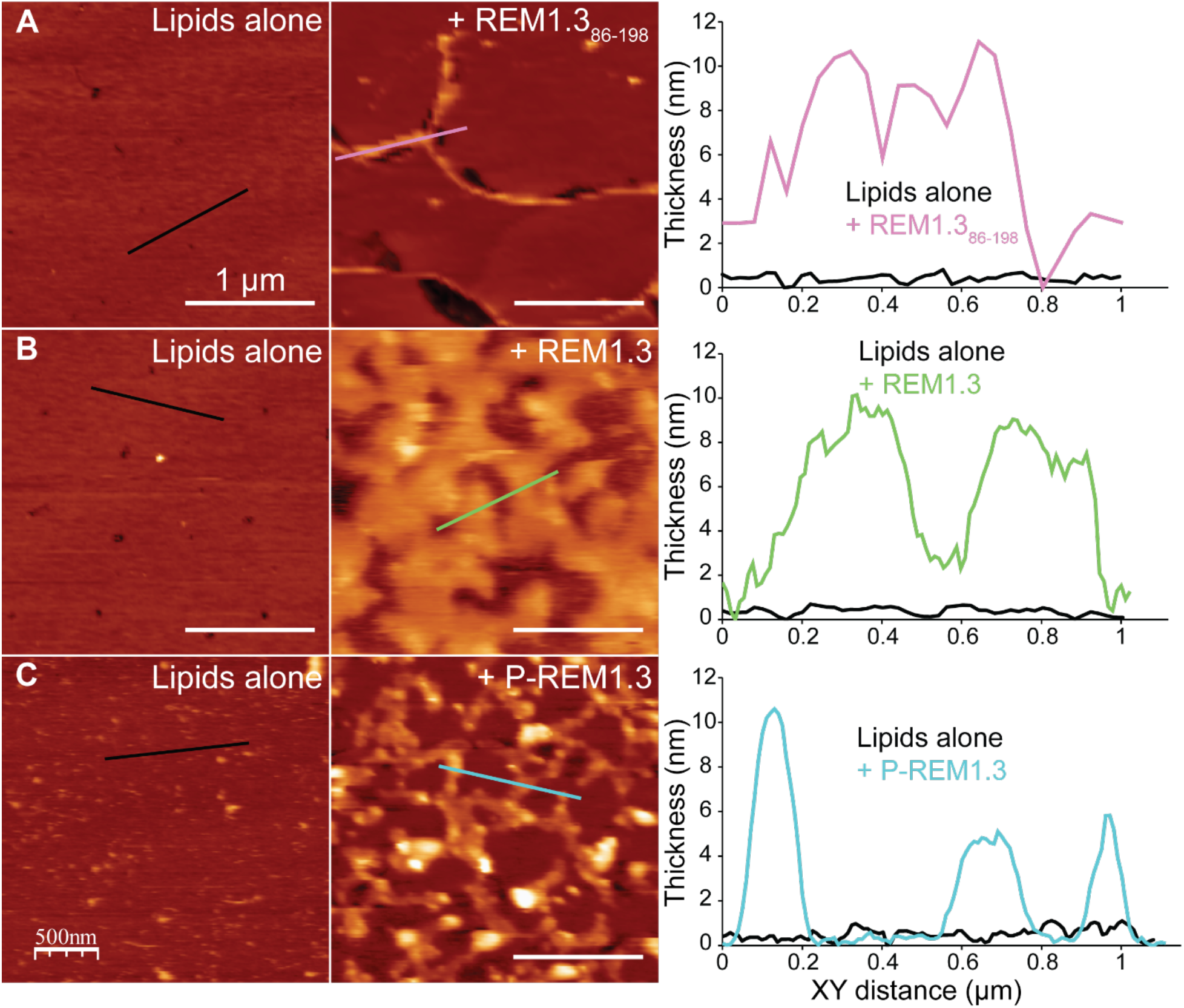
REM1.3-driven nanodomain formation by AFM. Small Unilamellar Vesicles (SUVs) made of DPPC/DLPC/sitosterol/PIPmix (50/18/8/24 mol%) are deposited on a mica plate. (A) A smooth membrane (left) generated by incubating the lipid mixture overnight at 30Cº, was incubated for 30 min with REM1.3_86-198_ (middle). The profiles under the two lines in the images are indicated in the right panel. (B,C) Similarly, a smooth membrane (left) was incubated for 30 min with Full length REM1.3 (middle) either non-phosphorylated (B) or phosphorylated by AtCPK3 (C). The profiles under the two lines in the images are indicated in the right panel. Images shown are representative of those found in at least 3 independent experiments. Scale bars: 1 μm.

We assessed whether pre-existing lipid domain would impact the formation of REM1.3 domains. We produced heterogeneous lipid bilayers with the same lipid composition as above, see Material and methods. In absence of protein, the membrane displayed two different heights, separated just by ≈ 1 nm, corresponding to two distinct lipid domains. Assuming that high-order lipid domains would be thicker than low-order ones (Grélard et al., 2013) and considering the greater size of PIPs molecules compared to PCs, we hypothesized that the highest domains were formed by PIPs and PS. After the addition of purified full length REM1.3, we observed the formation of protein domain of ca. 7 nm thick (Figure S4) and a restructuring of the existing lipid domains. This can be interpreted as indicating that REM1.3 is interacting with and redistributing the PIPmix-enriched domains. We further used the heterogeneously expressed Full length REM1.3 (purified without the GFP tag, Figure S1) to test the potential role of the IDD in domain formation and stability. AFM observations showed that it also promoted the formation of larger, patch-like domains with a thickness around 4-10 nm (Figure 5B).

### The shape and size of REM1.3 nanodomains are phosphorylation-dependent

REM1.3 can be phosphorylated by AtCPK3 *in vitro*, and the protein’s mobility in the membrane and nanodomain size are affected by its phosphorylation state *in vivo* (Perraki et al., 2018). Thus, we tried to recreate and analyze this behavior in our minimal *in vitro* system. We phosphorylated *in vitro* REM1.3 with AtCPK3 purified form *E. coli* (Figure S5).

The addition of phosphorylated REM1.3 to smooth membranes resulted in the creation of domains of 4-10 nm in thickness but much smaller width compared to non-phosphorylated REM1.3 (Figure 5C). In the case of non-phosphorylated REM1.3, the domains reached lateral dimensions of up to several hundred nanometers in width, reaching up to 1 μm, whereas the interconnected domains formed by the phosphorylated protein were not wider than 100-200 nm. This difference in phenotype *in vitro* is reminiscent of what has been observed *in vivo* by Single-Particle Tracking Photoactivated Localization (Gronnier et al., 2017; Perraki et al., 2018).

To summarize, AFM studies showed the ability of both truncated and Full length REM1.3 to trigger domain formation even in absence of pre-formed lipid domain in a smooth, homogeneous membrane. This implies that domain formation is partly protein-driven. The difference in organization pattern when lipid phase separations are pre-formed (thread-like for the truncated REM1.3_86-198_ vs. patch-like for the full length REM1.3) implies that, the IDD, absent in the truncated REM1.3_86-198_, must indirectly play a role in supramolecular membrane nanodomain organization even though the two proteins share the same membrane binding mechanism. Phosphorylations in the IDD of REM1.3 led to the disorganization of the membrane domains into smaller and more disperse domains.

## Discussion

The remorin proteins are frequently involved in signaling to regulate plant biotic and abiotic interaction with their environment (Gouguet et al., 2020). As such, Remorin’s lateral segregation to specific nanoscale compartments at the PM are issues under high scrutiny in the fields of plant biochemistry, biophysics and signaling.

### The set of minimal partners to form REM1.3 nanodomains: Driving forces behind REM1.3 nanodomain formation

In this study, we assessed whether a set of 3 classes of lipids was sufficient to reconstitute group 1 REM nanodomains *in vitro*: 1/ a mix of saturated and unsatured bilayer-forming PC to create lo and ld domains; 2/ negatively charged polyphosphoinositides that allow the loading of REM1.3 to the bilayer (Figure 2); 3/ phytosterols that are likely necessary to stabilize the nanodomains (Figure S3). All of these molecules are necessary to form the protein-lipid nanodomains located exclusively along the exposed leaflet of the bilayer where REM1.3 is organized in a periodic manner (Figure 4). Homomeric REM1.3 (Martinez et al., 2018) targets PI4P of the PM’s inner leaflet *in vivo*, and other anionic lipids to a lesser extent (Figure 2). This binding involves the RemCA domains through electrostatic interactions with the phosphate group of PIPs and the Lys/Arg of REM-CA (Gouguet et al., 2020; Gronnier et al., 2017) (Legrand et al., 2019), while sitosterol would likely migrate to the saturated fatty acid-enriched nascent anchoring site (Legrand et al., 2019). Finally, nearby REM1.3 would cluster together through their coiled-coil domains (Figure 5A) and more efficiently through the IDD (Figure 5B), increasing nanodomain size. Indeed, disruption of the coiled-coil homomerization domain, as exemplified by GFP-REM1.3_EEE_ greatly reduces, if not abolishes, membrane targeting (Figure 1). We envision that high-order REM1.3 interaction through the coiled-coil and IDD likely cluster more PIPs and sitosterol. This may act as a self-amplifying system driving the formation of nanodomains.

Size limitation of membrane nanodomains could come from the exhaustion of nanodomain components, particularly REM1.3 and PIPs, in the immediate vicinity of the nanodomain. We assume all nanodomain components are in equilibrium between two states: either in the bulk membrane or in membrane nanodomains. This implies nanodomain dissociation may overcome nanodomain formation once a certain nanodomain size is reached. To decipher such mechanisms would require a thorough kinetic analysis of nanodomain formation, which is beyond the scope of this study.

Whereas disruption of the coiled-coil homotrimerization domain abolishes membrane targeting, REM-CA alone still is capable of PM binding (Legrand et al., 2019). This underlines the cooperative interactions of the trimerization domain towards membrane association. Our previous observation of coiled-coil domain clustering in absence of membranes underlines the tendency of coiled-coil domain interactions.

Altogether, we conclude that reconstituting nanodomains of REM1.3 requires the following minimal set of partners: REM1.3_86-198_, i.e. RemCA domains bundled through a homomeric coiled-coil domain with 3 IDD (Martinez et al., 2018), PIPs, sitosterol (Gronnier et al., 2017; Legrand et al., 2019; Raffaele et al., 2009) and bulk lipids, such as DPPC and DLPC, to favor the formation of two distinct lipid phases, lo and ld respectively.

This leaves the question: what drives nanodomain formation? PIPs are known to segregate from the bulk of the membrane into nanodomains without requiring any protein or sterol (Bilkova et al., 2017; van den Bogaart et al., 2011; Furt et al., 2010; Ji et al., 2015; Motegi et al., 2021). Some sterols, like sitosterol, are also enriched in nanodomains in PC/PIP membranes (Furt et al., 2010). Lipid phase separation on supported bilayer for AFM can be achieved without REM1.3 (Figure S4). Thus, we wished to assess whether REM1.3 had an active role in shaping membrane nanodomains or if it only targeted pre-existing nanodomains. Variations in lipid domain surface occupancy by AFM upon addition of REM1.3 (Figure 5) with or without pre-existing formed lipid domains prove REM1.3’s ability to deeply modify membrane organization by forming 6-7 nm tall nanodomains. Thus, REM1.3 nanodomain formation is mostly protein-driven with the involvement of lipid to stabilize the REM domain by a feedback mechanism between proteins and lipids, see model proposed in Figure 6A.

**Figure 6.**
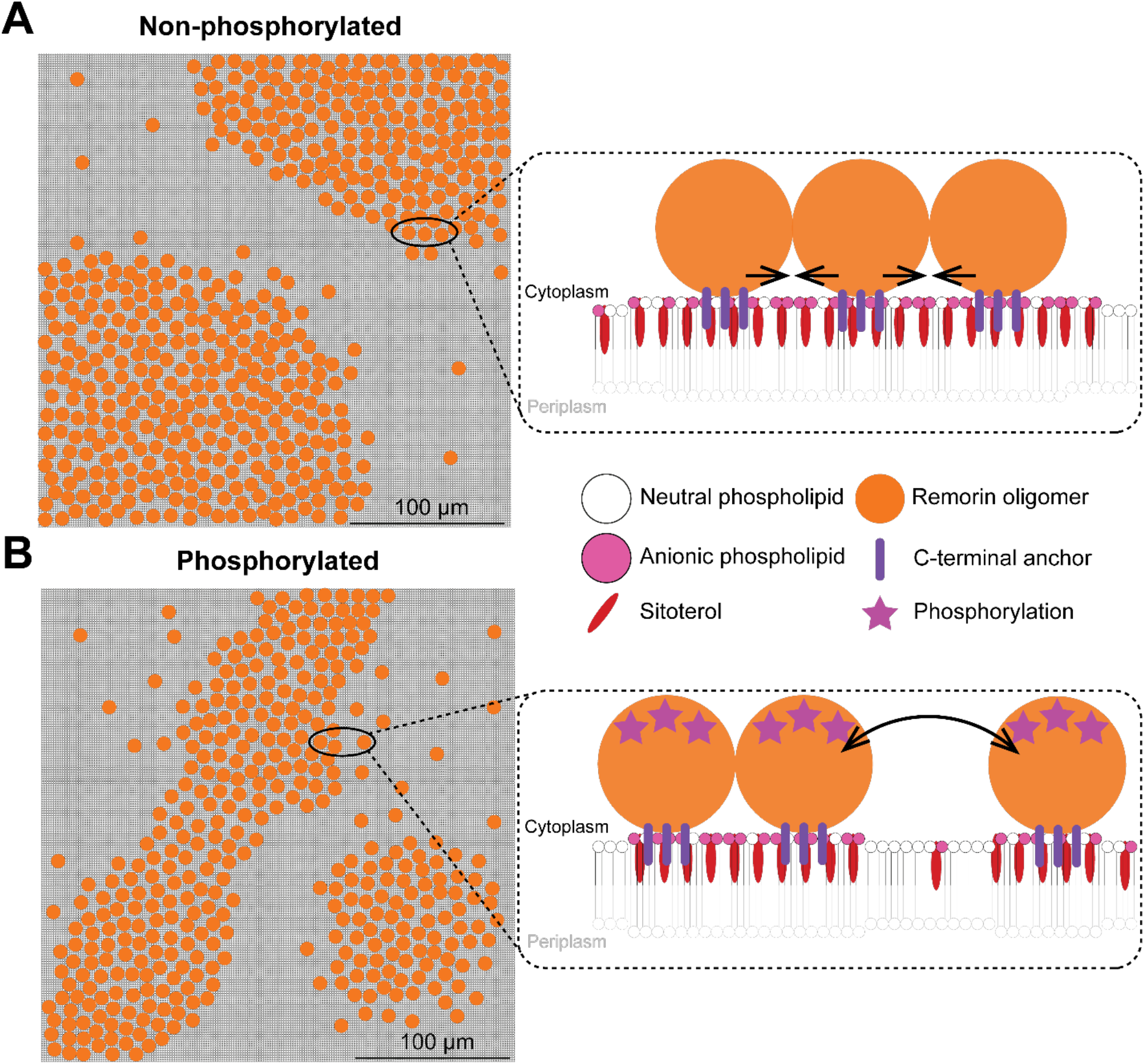
Schematic of the phospho-dependent REM1.3 membrane nanodomain formation. REM1.3 oligomers bind to anionic phospholipids (PIPs and PA but not PS) of the PM’s inner leaflet and cluster into membrane nanodomains in presence of sitosterol. Membrane nanodomain size depends on REM1.3’s phosphorylation state: (A) a few hundreds of nanometers if REM1.3 is not phosphorylated but (B) only about 100 nm if REM1.3 is phosphorylated. Insert: a close-up lateral view of this process depicts REM1.3 clustering with anionic phospholipids (PIPS and PA but not PS) and sitosterol, resulting in a slight increase in lipid order (Legrand et al., 2019). Protein-protein interactions (black arrows) and phosphorylations define the size and the microscopic organization pattern of REM1.3 membrane nanodomains.

Our data delineating both an impact of the coiled-coil oligomerization as well as of the multivalent electrostatic interactions in the IDD upon nanodomain clustering could suggest the creation of a membrane-associated liquid phase separation (Ditlev, 2021). Liquid-liquid phase separation (LLPS) is a longstanding concept that gains in importance through its role in multiple cellular processes (Li et al., 2012). IDD and polymerization domains, such as coiled-coil domains, are typically overrepresented in LLPS forming proteins (Alberti et al., 2019). Shaping the features of membrane targeting and nanodomain clustering by both domains demonstrates the role of the multivalent interactions that could engender phase separation on the membrane surface. The formation of a lo phase in the membrane could be orchestrated to the simultaneous creation of a separated liquid phase.

### Orientation of REM1.3 at the surface of the lipid bilayer

Different models were proposed to explain the spatial organization of group1 REM: either the coiled-coil is oriented perpendicular to the bilayer (Gouguet et al., 2020) or it lays at the surface of the membrane (Jaillais and Ott, 2020). The estimated length of the coiled-coil being ca. 8 nm with a diameter of 2 nm (Martinez et al., 2018) is compatible with the height observed with the AFM on the supported lipid bilayers, but not with the cryo-EM observation of GFP-REM1.3_86-198_ protruding at only 4.5 nm on the surface. This would rather suggest an orientation of REM1.3 with its coiled-coil laying at the surface of the bilayer, with the REM-CAs embedded in the membrane like staples, and the IDDs interacting together. Although this hypothesis remains to be experimentally proved, it paves the ways towards elucidating the precise orientation of REM1.3 in membrane nanodomains.

### Phosphorylation of REM1.3 affects nanodomain organization

The phosphorylation of REM1.3 by AtCPK3 yields a protein, P-REM1.3, that triggers a different nanodomain organization than the non-phosphorylated REM1.3 (Figure 5C). P-REM1.3 nanodomains have the same height but display a less compact and more disperse lateral packing on the surface of the bilayer. This data is in line with live super-resolution single particle tracking microscopy where we observed that the phosphorylation increases the EOS-REM1.3’s diffusion coefficient and mean square displacement reflecting an increase of REM1.3 mobility (Perraki et al., 2018). Moreover, Voronoï tessellation method that compare the *in vivo* supra-molecular organization of microscopy data showed a decrease in the localization density of EOS-REM1.3 in nanodomains after phosphorylation, i.e. after infection by the Potato Virus X (Perraki et al., 2018). Overall, the changes of REM1.3 distributions in synthetic membranes after phosphorylation phenocopy the increase of REM1.3’s modulation of nanodomain organization. This data suggests the involvement of the phosphorylation of the IDD in the organization of the protein in the lipid bilayer, likely inducing conformational changes that would hinder REM1.3 oligomerization through an uncharacterized mechanism. The phosphorylated IDD could act as an entropic barrier to control nanodomain size, effectively pushing away other REM1.3 homotrimers and hindering nanodomain formation (Jamecna et al., 2019). Alternatively, the phosphorylation could also modulate electrostatic interactions and repulsions with other IDDs (Liu et al., 2014). Regulation by phosphorylation is also in line with our previously mentioned suggestion of LLPS formation on the membrane surface, since phosphorylation is involved in LLPS modulation events (Wang et al., 2021).

Our present work of *in vitro* reconstitution paves the way to understand the labelling of different REM phylogenic group that localize to different nanodomains and are in some cases completely excluded from each other (Jarsch et al., 2014).

## Supporting information

Supplemental informations

## Acknowledgement

We thank the Bordeaux Imaging Center, part of the National Infrastructure France-BioImaging supported by the French National Research Agency (ANR-10-INBS-04). We thank Bordeaux-Metabolome platform for lipid analysis (https://www.biomemb.cnrs.fr/en/lipidomic-plateform/) supported by Bordeaux Metabolome Facility-MetaboHUB (grant no. ANR–11–INBS–0010 to SM). This work was supported by the French National Research Agency (grant no. ANR-19-CE13-0021 to SM, VG, MB), the European Research Council (ERC) under the European Union’s Horizon 2020 research and innovation program (Grant Agreement No 101001097 to YJ). The IPS2 benefits from the support of the LabEx Saclay Plant Sciences-SPS (ANR-10-LABX-0040-SPS). We thank Frederic Daste for his help in the protocole of GUV by the Teflon method.

The authors declare no conflict of interest

## Author’s contribution

ALe purified the proteins and performed the GUV experiments; G-C performed and analyzed the AFM; MDJ measured the clustering of REM1.3 in pss1 and WT; MD, OL performed and analyzed cryo-EM experiments; VB and YJ produced the pss1 line expressing fluorescent REM1.3; MB and ALe performed the phosphorylation experiments of REM1.3; BH, VG and ALo advised on biophysical analyses; ALe, MV and SM designed the project, analyzed the data, built the figures and wrote the manuscript.

## Supplementary information

**Figure S1. Purity of proteins used in this study**.

GFP-REM1.3_86-198,_ GFP-REM1.3_86-198, EEE_, REM1.3_86-198_, and REM1.3 were loaded on a tris-tricine 13% (w/v) polyacrylamide gel. Protein standard (ladder) was Mark 12 protein standard from ThermoFisher. Stars indicate GFP-REM1.3_86-198_ and GFP-REM1.3_86-198,EEE_, respectively.

**Figure S2. GFP-REM1.386-198 binds to GUVs enriched with 32 mol% POPS**.

A GUV containing DPPC/DLPC/sitosterol/POPS 40/21/6/32 mol% is incubated with GFP-REM1.3_86-198_. *Left*: RhodPE channel (magenta). *Middle*: GFP-REM1.3_86-198_ channel (green). *Right*: composite image from both channels. Scale bar: 5 μm. Observation statistics are given in Table S1.

**Figure S3. PIPs and β-sitosterol are required for nanodomain formation**.

A bilayer composed of DPPC/DLPC/PIPmix (50/18/24 mol%) (left) is incubated with 2 μM of REM1.3_86-198_ for 30 min (middle). Thickness profiles of lines drawn on both images are provided (right).

**Figure S4. Full-length REM1.3 forms nanodomains in a supported bilayer with pre-existing lipid domains**

AFM observation of a patchy membrane with pre-existing lipid domain (generated without heating the lipid mixture) was incubated for 30 min with Full-length REM1.3. Two thickness representative profiles are provided (right). Experiments were performed 4 times with different mica substrates, and representative are presented in the Figure. Scale bars: 1 μm.

**Figure S5. *In vitro* phosphorylation assay of REM1.3 by AtCPK3**

6His-REM1.3 and GST-AtCPK3 (WT and dead D202A) were purified from *E. coli* and mixed to a ratio of 4:1 for *in vitro* kinase assay. Bands corresponding to autophosphorylation of GST-AtCPK3 and transphosphorylation of 6His-REM1.3 are indicated (top). Gel was stained by coomassie blue to visualize protein loading (bottom).

**Table S1. Statistics for GUV observations by confocal microscopy**.

Number of observed GUVs (observed liposomes) and number of GUVs showing GFP-REM1.3_86-198_ binding (binding events). Binding frequency: ratio of the number of the number of binding events over the number of observed liposomes. ϕ: DPPC/DLPC/sitosterol 50/42/8 mol% GUVs in presence of GFP-REM1.3_86-198_. PIPmix _(EEE)_: DPPC/DLPC/sitosterol/PIPmix 50/26/8/16 mol% GUVs in presence of GFP-REM1.3_86-198,EEE_ (Figure 1B). High PS: DPPC/DLPC/sitosterol/POPS 40/21/6/32 mol% GUVs in presence of GFP-REM1.3_86-198_ (see Figure S2).

