## Supplemental informations for "Structural determinants of REMORIN nanodomain formation in anionic membranes"

**Figure S1**

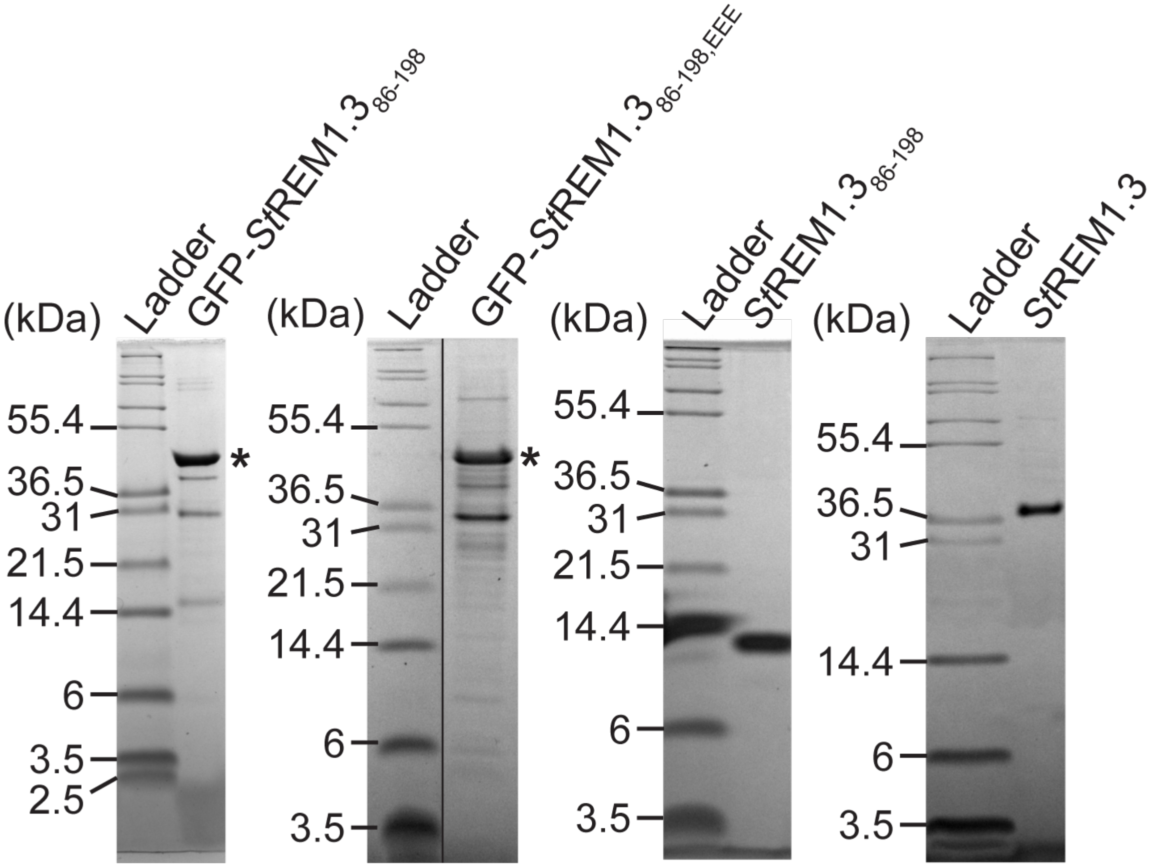


**Figure S1. Purity of proteins used in this study.**
GFP-REM1.3$_{86\text{-}198}$, GFP-REM1.3$_{86\text{-}198,\ EEE}$, REM1.3$_{86\text{-}198}$, and REM1.3 were loaded on a tris-tricine 13% (w/v) polyacrylamide gel. Protein standard (ladder) was Mark 12 protein standard from ThermoFisher. Stars indicate GFP-REM1.3$_{86\text{-}198}$ and GFP-REM1.3$_{86\text{-}198,EEE}$, respectively.

**Figure S2**

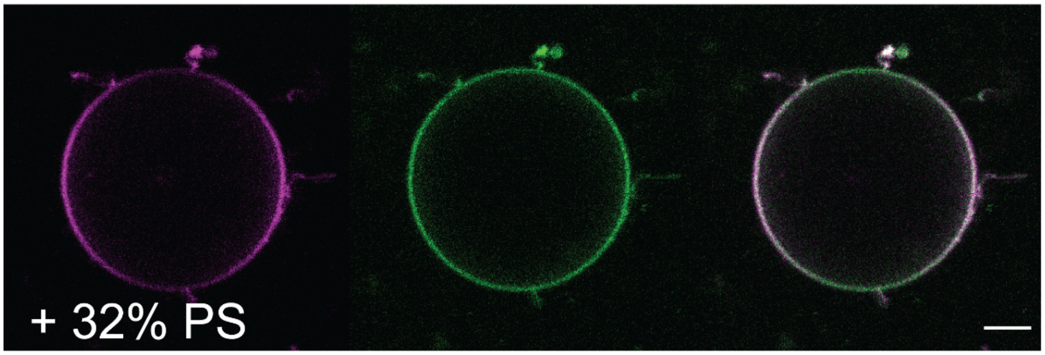


**Figure S2. GFP-REM1.386-198 binds to GUVs enriched with 32 mol% POPS.**
A GUV containing DPPC/DLPC/sitosterol/POPS 40/21/6/32 mol% is incubated with GFP-REM1.3$_{86-198}$. *Left*: RhodPE channel (magenta). *Middle*: GFP-REM1.3$_{86-198}$ channel (green). *Right*: composite image from both channels. Scale bar: 5 µm. Observation statistics are given in Table S1.

**Figure S3**

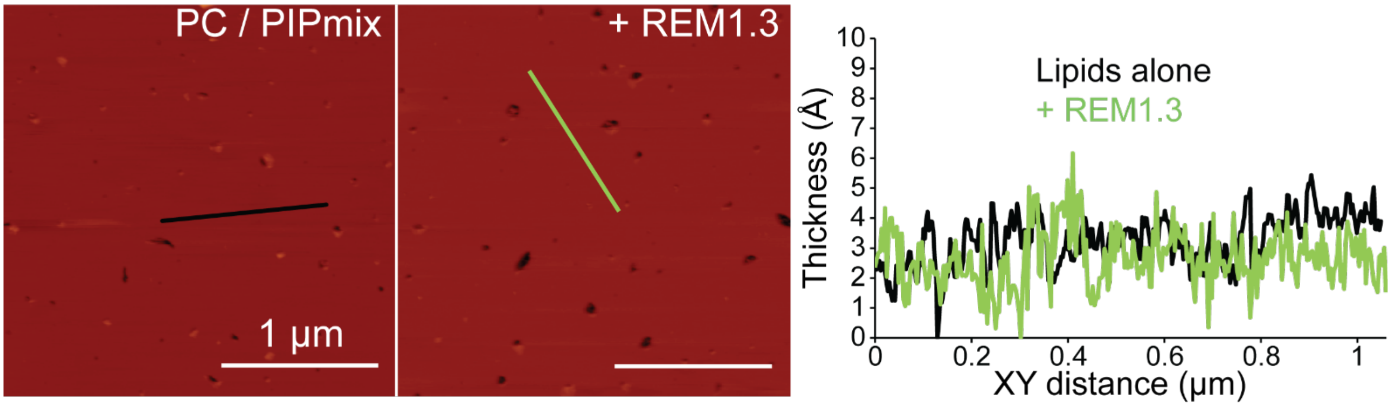


**Figure S3. PIPs and β-sitosterol are required for nanodomain formation**.
A bilayer composed of DPPC/DLPC/PIPmix (50/18/24 mol%) (left) is incubated with 2 µM of REM1.3$_{86\text{-}198}$ for 30 min (middle). Thickness profiles of lines drawn on both images are provided (right).

**Figure S4**

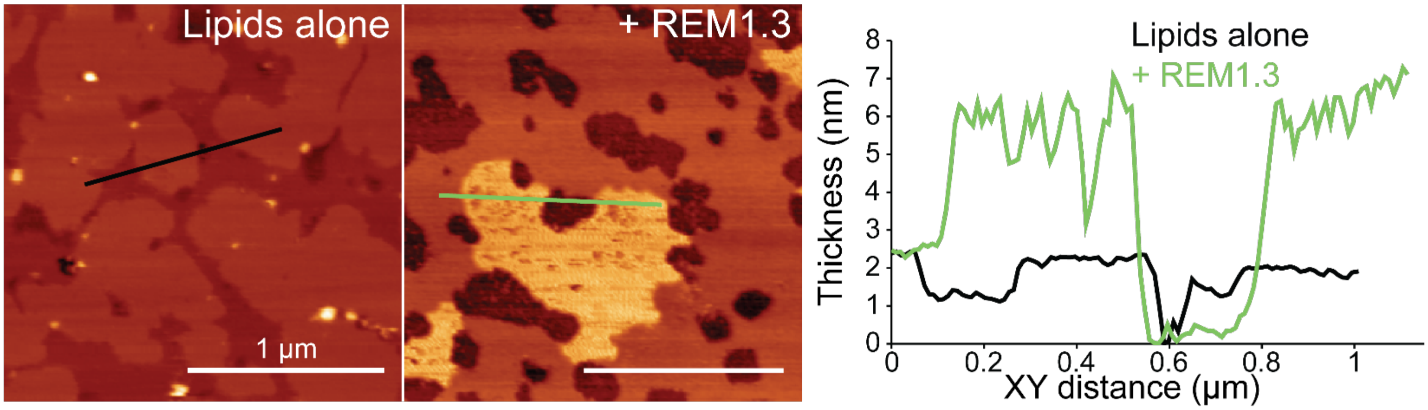


**Figure S4. Full-length REM1.3 forms nanodomains in a supported bilayer with pre-existing lipid domains**

AFM observation of a patchy membrane with pre-existing lipid domain (generated without heating the lipid mixture) was incubated for 30 min with Full-length REM1.3. Two thickness representative profiles are provided (right). Experiments were performed 4 times with different mica substrates, and representative are presented in the Figure. Scale bars: 1 µm.

**Figure S5**

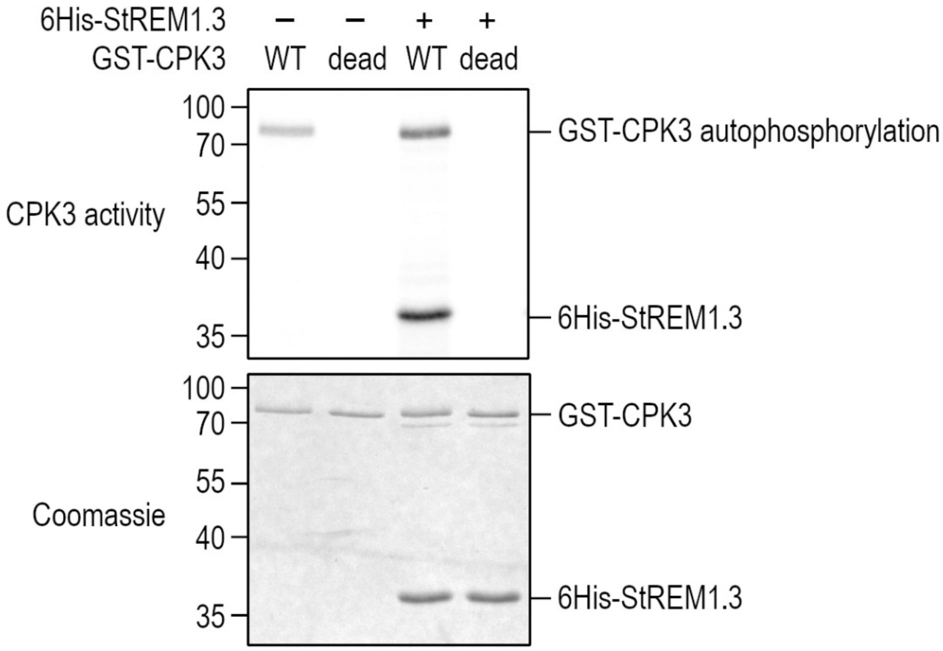


**Figure S5. *In vitro* phosphorylation assay of REM1.3 by AtCPK3**
6His-REM1.3 and GST-AtCPK3 (WT and dead D202A) were purified from *E. coli* and mixed to a ratio of 4:1 for *in vitro* kinase assay. Bands corresponding to autophosphorylation of GST-AtCPK3 and transphosphorylation of 6His-REM1.3 are indicated (top). Gel was stained by coomassie blue to visualize protein loading (bottom).

**Table S1. Statistics for GUV observations by confocal microscopy.**

Number of observed GUVs (observed liposomes) and number of GUVs showing GFP-REM1.3$_{86-198}$ binding (binding events). Binding frequency: ratio of the number of the number of binding events over the number of observed liposomes. ϕ: DPPC/DLPC/sitosterol 50/42/8 mol% GUVs in presence of GFP-REM1.3$_{86-198}$. PIPmix $_{(EEE)}$: DPPC/DLPC/sitosterol/PIPmix 50/26/8/16 mol% GUVs in presence of GFP-REM1.3$_{86-198,EEE}$ (Figure 1B). High PS: DPPC/DLPC/sitosterol/POPS 40/21/6/32 mol% GUVs in presence of GFP-REM1.3$_{86-198}$ (see Figure S2).

| Condition | Observed liposomes | Binding events | Binding frequency |
|---|---|---|---|
| ϕ | 17 | 0 | 0 |
| PIPmix | 374 | 287 | 0.767379679 |
| PIPmix (EEE) | 205 | 12 | 0.058536585 |
| POPI4P | 187 | 103 | 0.550802139 |
| POPI5P | 1156 | 385 | 0.333044983 |
| DOPI(4,5)$P_2$ | 327 | 208 | 0.636085627 |
| PAPI(3,4,5)$P_3$ | 17 | 16 | 0.941176471 |
| 16% POPS | 272 | 0 | 0 |
| 32% POPS | 311 | 195 | 0.627009646 |
| POPA | 974 | 134 | 0.137577002 |
| GIPC | 5 | 0 | 0 |
